# Unlocking the potential of *miR-19b* in the regulation of temozolomide response in glioblastoma patients via targeting PPP2R5E, a subunit of the protein phosphatase 2A complex

**DOI:** 10.1101/2023.01.16.524069

**Authors:** Elham Kashani, Kristyna Hlavackova, Stefan Haemmig, Martin C Sadowski, Jaison Phour, Ulrich Baumgartner, Nicole Mueller-Wirth, Carmen Trefny, Bushra Sharf Den Abu Fakher, Coline Nydegger, Theoni Maragkou, Philippe Schucht, Aurel Perren, Pascal Zinn, Markus Lüdi, Thomas Michael Marti, Philippe Krebs, Erik Vassella

**Author notes:** Corresponding author: Prof. Dr. Erik Vassella Institut für Gewebemedizin und Pathologie, Murtenstrasse 31, 3008 Bern, Switzerland.

## Abstract

Despite the standard of care, glioblastoma IDH wildtype (GBM) inevitably recurs, underscoring the need to develop new treatment strategies. To address the role of microRNAs in temozolomide (TMZ) response, we performed functional microRNA screens and consistently identified *miR-19b*. Our study reveals a novel axis between *miR-19b* and PPP2R5E subunit of serine/threonine protein phosphatase PP2A and establishes a so far unappreciated contribution of *miR-19b* in TMZ resistance of GBM. Specifically, our results demonstrate that attenuation of *miR-19b* in GBM cell lines and glioblastoma stem cells (GSCs) induces DNA damage, which further enhances the cytotoxic effects of TMZ treatment. We confirmed TMZ resistance induced by knocking down PPP2R5E in orthotopic mouse xenografts of GSCs. Furthermore, our results indicate that treating cells with the PP2A-activating drug FTY720 or knocking down endogenous PP2A-inhibiting proteins potentiates the cytotoxic effects of TMZ. *MiR-19b* attenuation or PPP2R5E activation could potentially be exploited in adjuvant therapy of GBM patients.

## Background

Glioblastoma, IDH wildtype (GBM), the most common and most aggressive primary brain tumor in adults, remains an oncology challenge with a median two-year survival rate of only 3.3% ^1^. The current standard of care for patients diagnosed with GBM consists of a combination therapy regimen, which includes surgical resection, treatment with the DNA alkylating agent temozolomide (TMZ), and radiation therapy. Administration of TMZ results in the formation of cytotoxic DNA adducts, including O^6^-methylguanine (O^6^-MeG), N^7^-methylguanine and N^3^-methyladenine ^2^. If left unrepaired, these adducts may trigger senescence and apoptosis. O^6^-MeG alkylation can be repaired in a single-step reaction mediated by O^6^-MeG DNA methyl transferase (MGMT), leading to poor TMZ response. However, in 45% of GBM patients, MGMT is silenced by promoter hypermethylation, which is associated with a better therapy response ^3^. If left unrepaired, O^6^-MeG will mispair with thymine, leading to futile cycles of mismatch repair (MMR). This can promote the occurrence of one-ended DNA double stranded breaks (DSBs) owing to collapsed replication forks during DNA replication. Ataxia telangiectasia mutated (ATM) and ataxia telangiectasia Rad3-related (ATR) are activated in response to DNA DSBs, initiating a cascade activation of downstream effectors, including H2A.X variant histone (H2AX) and checkpoint kinases CHK1 and CHK2. This cascade ultimately induces cell cycle arrest in the G2/M phase and promotes DNA repair ^4^. Notably, glioblastoma stem cells (GSCs) exhibit intrinsic resistance to radio-chemotherapy, which is attributed to their resistance to apoptosis and increased DNA repair capacity ^5^.

Acquired resistance to TMZ may be due to an increase in the expression of MGMT ^6^ or may alternatively be caused by acquisition of loss-of-function mutations or downregulation of mismatch repair genes ^7^. Enhanced homologous recombination activity may be an alternative mechanism of acquired TMZ resistance ^4,8^. Activation of anti-apoptotic pathways, autophagy and senescence are additional survival strategies of GBM cells to cope with TMZ-induced cytotoxic effects ^2^. Longitudinal analyses of recurrent GBM were performed to identify additional alterations that drive TMZ resistance ^9,10^. However, there was no evidence of persistent secondary alterations in the recurrent tumors. Instead, high plasticity was identified as the primary cause of tumor recurrence ^9^.

MicroRNAs (miRNAs) are critical post-transcriptional regulators of gene expression that are also involved in DNA repair pathways, contributing to a fine-tuning regulation of DNA repair or survival pathways (for review, see ^11^). However, their role in TMZ resistance has yet to be systematically analyzed.

In this study, we performed unbiased systematic screening for miRNAs that confer TMZ resistance in GBM and reproducibly identified cells that overexpress *miR-19b*. We showed that *miR-19b* targets the PPP2R5E regulatory subunit of serine/threonine protein phosphatase PP2A, thereby uncovering a new mechanism by which *miR-19b* controls TMZ resistance in GBM.

## Methods

### GBM patient cohorts

Two retrospective GBM cohorts were used in this study: the miRNA screen discovery cohort (n=28), consisting of therapy-naive IDH wt GBM ^12^, and the longitudinal cohort (n=43), consisting of matched initial (therapy-naïve) and recurrent tumors with an interval of at least one year between two resections. Ten normal adjacent brain tissue samples were also included in the latter cohort ^10^.

### Cell lines and primary cells

Human primary glioblastoma stem cells (GSC3 and GSC4) were collected intraoperatively from patients with GBM IDH wt at MD Anderson ^13^. The GSCs were cultured in GSC media (DMEM/F12 containing 1× B27, 20 ng/ml EGF, and 20 ng/ml basic fibroblast growth factor). The mutational profile of GSCs was assessed using the TruSight Oncology 500 cancer panel (Illumina) and a NovaSeq 6000 instrument. Identified mutations are shown in suppl.Table1.

U87MG (kindly provided by M.E Hegi, University of Lausanne, Switzerland), LN18 and LN229 (purchased from American Type Culture Collection) were cultured with 10% fetal bovine serum (Sigma) at 37°C and 5% CO_2_ ^12^ at maximally 70% confluence. Cells were genotyped and authenticated by Microsynth (Switzerland) and tested negative for mycoplasma infection. Unless otherwise stated, cells were treated with [100 μM] TMZ (Selleck Chemicals).

### Patient-derived xenograft (PDX) mouse model

Groups of 5-week-old nude (athymic (Crl:NU(NCr)-Foxl nu)) mice (Charles River) were anesthetized and orthotopically injected into the right forebrain with 1-2×10^5^ GSC4 cells co-transduced with a beetle luciferase reporter and sh(ctrl) or shPPP2R5E constructs ^14^. All animals were treated with a dosage of 66 mg/kg TMZ by oral gavage daily for five consecutive days starting two weeks after engraftment and then rested for 3 weeks. This procedure was repeated two more times for a total of three cycles. Bioluminescence was used to longitudinally monitor tumor size, which was measured by an *in vivo* imaging system (SpectrumCT In Vivo Imaging System (IVIS), PerkinElmer, MA, US). Brain and spine signals intensities were analyzed by Living Image® software (PerkinElmer, MA, US) ^15^.

Mice were euthanized at the onset of neurological symptoms or upon severe weight loss.

### Constructs, transfections, and transductions

Luciferase reporter plasmids were constructed using the pmiRGLO Dual-Luciferase miRNA Target Expression Vector (Promega, Dübendorf, Switzerland) ^16^. Generation of lentiviruses and lentiviral transduction of GBM cell lines were performed as described ^16^. Transient transfection of short RNAs was performed as described ^12^. The sequences of oligonucleotides and lentiviral constructs used in this study are indicated in suppl.Table2.

### MiRNA lentivirus screen and next generation sequencing (NGS)

2.5×10^6^ IFU/ml of pooled lenti-miRNA lentivirus library (System Biosciences Cat. #PMIRHPLVHT-1) was mixed with 4 mg/mL polybrene (Sigma) and transduced into 5×10^5^ cells at a MOI of 0.3-0.4 to ensure single lenti-miR integration. Three days post-transduction, cells were treated with 1.5 μg/ml puromycin (Sigma) for three days and subsequently treated with TMZ. Selection was started with 25 μM TMZ, and increased to 100 μM.

For NGS analysis, lentiviral fragments encompassing the pre-miRNA sequence were amplified by PCR using the High-Fidelity MasterMix (NEB) and 400 ng genomic DNA, 0.5 μM primers and 1 mM MgCl_2_ for 30 cycles. DNA fragmentation was performed using the Next Fast DNA Fragmentation & Library Prep Set (NEB). 100 ng purified DNA library was ligated with 2 μl barcode and 2 μl P1 (ThermoFisher). NGS analysis was performed using an Ion Torrent PGM (ThermoFisher). Differential expression analysis was performed using the RNA Sequencing Data Analysis Workflow from CLC Genomics Workbench (Qiagen). Relative coverage was calculated to the total reads.

The fragment size distribution during TMZ selection was assessed using a 6-carboxyfluorescein-labeled reverse primer in a PCR assay and a GA3500 genetic analyzer (ThermoFisher).

### MRNA analysis

RNA extraction and Reverse Transcription quantitative real-time PCR (RTqPCR) was performed as described ^16^. Primers used for RTqPCR are shown in suppl.Table1.

Microarray gene expression profiling using NanoString cancer progression custom panel and bioinformatic analysis was performed as described ^10^.

### Cell based assays

Luciferase reporter assays, cell viability assays using alamarBlue and Resazurin, caspase3/7 activity using the ApoTox-Glo Triplex kit (Promega), BrdU incorporation assay, and colony formation assay were performed essentially as described ^16^. For colony formation assay cells were treated with 15-30 μM TMZ for two weeks initiating 16h post-seeding.

For the β-galatosidase (SA-β-gal) staining assay, 3×10^4^ cells were seeded in 6-well plates and treated with 100 μM TMZ for 48h, starting 16h post-seeding. SA-β-gal assay was performed using the senescence β-galactosidase staining kit (Cell Signaling #9860). At least ten randomly selected fields from the 20X magnification of the bright-field microscope (∼500 cells) were imaged and counted.

For γH2AX foci formation assay, 2000 cells were seeded in Ibidi optical 96-well plates (SLS4515-5EA). Cells were fixed with 2% PFA at room temperature for 10 minutes. Fixed cells were permeabilized using 0.5% Triton-X in PBS and blocked with 10% BSA/PBS for 1h at room temperature before incubation with primary anti-γH2AX antibody (1:500, CellSignaling#2577) overnight at 4°C. Cells were washed three times with DPBS and incubated with secondary Cy3-conjugated antibody (anti-rabbit). Images were acquired using an Incell Analyzer2000. Image analysis and foci quantification was performed using Cell Profiler (v.4.0.7).

### Western Blot

Western blots and antibodies are described elsewhere ^16^. Additional antibodies used in this study are anti-CIP2A (SantaCruz#sc-80662, 1:1000), anti-SET (SantaCruz# sc-133138,1:500), anti-phospho-Histone H2AX (Ser139) (cellSignaling#2577S, 1:1000), anti-PHAP1 (abcam#ab155148, 1:1000), anti-CHK1(Ser345) (CellSignaling,133D3, 2348, 1:1000), and anti-CHK2(Thr68) (CellSignaling,C13C1, 2197, 1:1000), and anti-PPP2CA (PP2A-Cα/β Antibody (1D6) HRP (LabForce,sc-80665 HRP,1:800)). Protein levels were normalized to total protein.

### Flow Cytometry

Cells were fixed with cold methanol and incubated with DAPI at room temperature for 10 minutes in the dark. Analysis of the cell cycle distribution was performed using a BD SORP LSR II flow cytometer connected to BD FACSDiva software. The data were analyzed using FlowJo V10 (Tree Star, Inc.(Ashland, OR, USA). The gating strategy involved eliminating dead cells and debris, and the G1 peak was used to adjust the S and G2/M peaks.

### TCGA data analysis

cBioPortal and TCGABiolinks R packages were used for survival data analysis. Overall survival was defined as the time interval between date of death or last contact and the date of initial surgery.

### Statistics

Statistical tests were performed using GraphPad Prism v.8.2.1 (GraphPad Software, San Diego, CA) and R studio v.4.0.3 (RStudio, Boston, MA). For comparing differences between two groups, non-parametric Mann-Whitney-U test was applied and p-values of <0.05 were considered as statistically significant. Unless otherwise stated, at least three biological replicates and three technical replicates were included in each experiment.

## Results

### miR-19b-3p overexpressing GBM cells are enriched by TMZ treatment

To identify miRNAs associated with TMZ resistance, U87MG and LN229 GBM cell lines were transduced with a pooled lentiviral expression library containing 520 miRNA precursors (pre-miRNAs) and were cultured at escalating doses of TMZ (suppl.Fig.1a). No viable cells were observed in the non-transduced group after TMZ treatment. Selection of resistant cells was monitored by PCR using primers encompassing pre-mRNA regions (suppl.Fig.1b, upper left). A discrete fragment pattern of PCR products, indicating selection, was observed 46 days post TMZ treatment (suppl.Fig.1b). NGS analysis showed a significant correlation of TMZ-enriched pre-miRNA sequences in two independent U87MG screens (suppl.Fig.1c-d). Comparison of the normalized coverage of individual pre-miRNA sequences revealed significant enrichment of *miR-19b* and *miR-20b*, which form part of the same pre-miRNA, in two independent U87MG screens and one LN229 screen (Fig.1a). In contrast, other pre-miRNAs were enriched exclusively in U87MG or LN229 cultures, respectively.

**Fig. 1.**
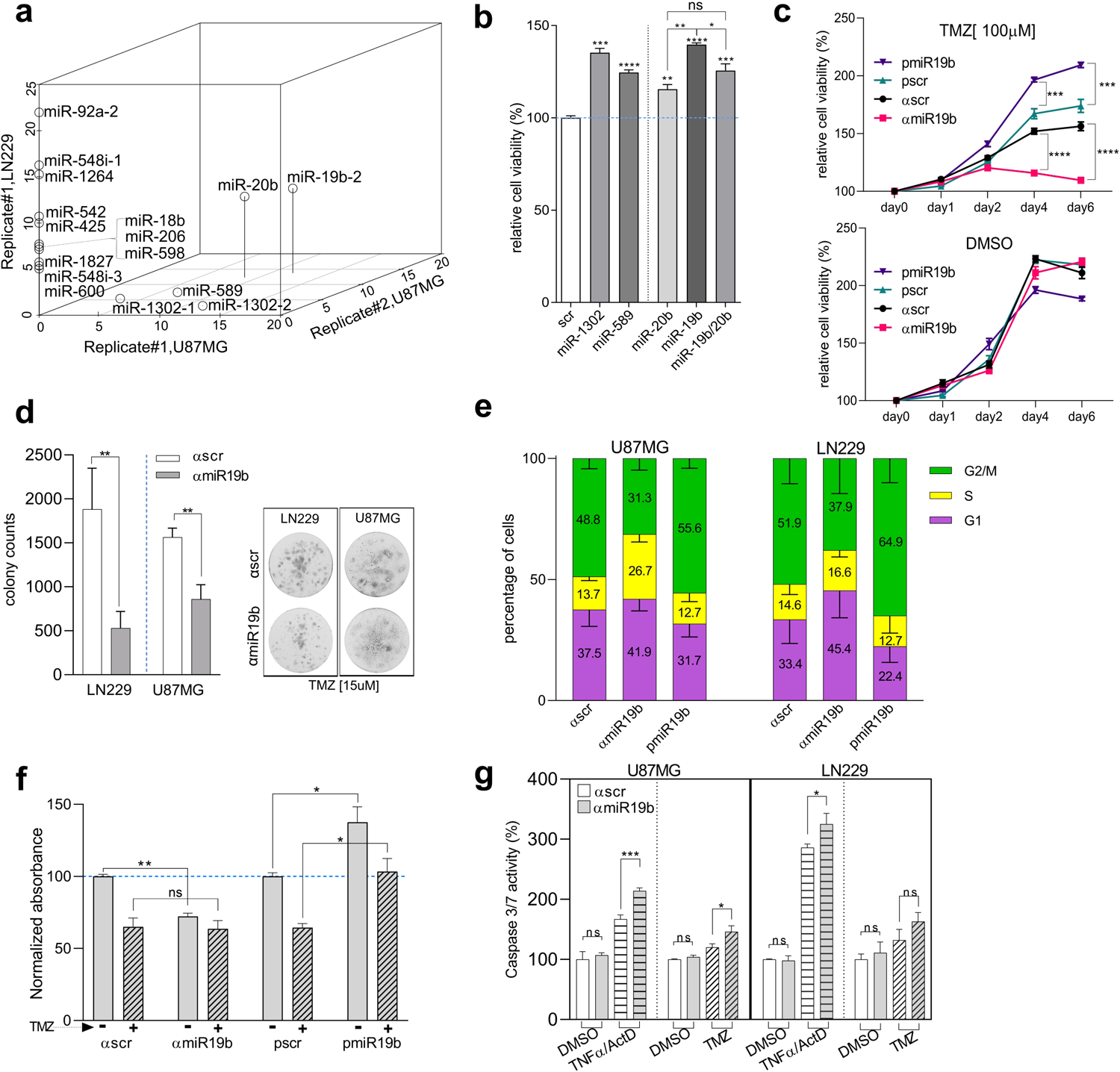
Systematic miRNA screen and cellular processes affected by *miR-19b*. **a** U87MG (x- and z-axis) and LN229 cells (y-axis) transduced with pooled pre-miRNA library and reproducibly enriched >5-fold following TMZ treatment relative to the DMSO control (p<0.028). **b** Cell viability by alamarBlue of U87MG cells transduced with pre-*miR-19b/20b*, *pre-miR-589*, and *pre-miR-1302* identified in (A) following TMZ treatment for 72h relative to pscr (n=3). **c** Cell viability by alamarBlue of U87MG cells transduced with miRNA constructs in presence of TMZ (upper) or DMSO control (lower). **d** Colony formation of U87MG cells in the presence of continuous treatment with 15 μM TMZ (n=3) and representative images of crystal violet-stained colonies (right) of U87MG and LN229 cells. **e** Cell cycle distribution following TMZ treatment (mean±SD, n=3). The percentage of cells in each phase is indicated in numbers. **f** BrdU incorporation assay of U87MG cells 4 days post treatment with TMZ (n=4). **g** Apoptosis induced with TNFα/ActD or 200 μM TMZ by caspase 3/7 activity assay.

To validate the results of the lentiviral screen, individual expression constructs for pre-miRNAs identified in the screens were used to transduce U87MG cells, which were then analyzed for TMZ response. Additionally, *pre-miR-19b-2* (pmiR19b) and *pre-miR-20b* were overexpressed individually in U87MG cells. In agreement with the screen data, all identified pre-miRs including *pre-miR-19b/20b*, *pre-miR-589*, and *pre-miR-1302* conferred resistance to TMZ (Fig.1b). Notably, pmiR19b alone conferred improved survival in the presence of TMZ compared to *pre-miR-20b* alone or *pre-miR-19b/20b*. In addition, *miR-19b* expression was associated with an unfavorable prognosis in an in-house “discovery” cohort of GBM IDH wt patients (n=28) (suppl.Fig.1e). We therefore focused on *miR-19b* for subsequent experiments.

### MiR-19b confers TMZ resistance in cell lines and GBM stem cells

We then undertook a series of tests to substantiate the contribution of *miR-19b* to TMZ resistance. Overexpression of pmiR19b resulted in enhanced survival of U87MG cells treated with TMZ, while overexpression of antisense *miR-19b* (αmiR19b) led to reduced survival compared to precursor scrambled control (pscr) or antisense scrambled control (αscr), respectively (Fig.1c, upper and suppl.Fig.2a). No difference in population growth was observed in DMSO control (Fig.1c, bottom). Consistently, clonogenic growth in the presence of TMZ was enhanced by pmiR19b (suppl.Fig.2b), whereas αmiR19b resulted in reduced clonogenic growth of U87MG or LN229 cells (Fig.1d).

We next assessed whether *miR-19b* expression correlates with stemness. Indeed, *miR-19b* levels were higher in GSC3 and GSC4, as well as U87MG which has retained stem cell features ^17^, compared to more differentiated LN229 and LN18 cells (suppl.Fig.2c). Attenuation of *miR-19b* in GSC4 also resulted in reduced viability in the presence of TMZ (suppl.Fig.2d). However, GSC3, which grows in large spheroids, gave rise to inconsistent results.

### Cellular processes elicited by miR-19b in response to TMZ

Following extensive dsDNA breaks, TMZ induces cell cycle arrest in G2/M after the next round of replication. When U87MG or LN229 cells were treated with TMZ for two days, G2/M arrest was observed in approximately 50% of the population (Fig.1e, αscr). However, attenuation or overexpression of *miR-19b* resulted in fewer (31-38%) or more (56-65%) cells in G2/M, respectively (Fig.1e), indicating that *miR-19b* enhances TMZ-induced G2/M arrest.

To assess the percentage of cells that were able to escape cell cycle arrest, we performed a 5-Bromo-2′-Deoxyuridine (BrdU) incorporation assay after the next round of replication during TMZ treatment. A higher proportion of cells overexpressing pmiR19b, compared to cells expressing pscr, were able to escape G2/M arrest in presence of TMZ, as indicated by a higher percentage of BrdU-positive cells, consistent with enhanced resistance (Fig.1f). In the absence of TMZ, pmiR19b cells showed increased BrdU incorporation, whereas cells with attenuated *miR-19b* expression showed reduced BrdU incorporation compared to the control (Fig.1f), in agreement with the oncogenic role of *miR-19b* ^16^.

Extensive accumulation of DNA damage leads to the induction of apoptosis ^2^. However, although TMZ-induced apoptosis was only slightly enhanced, the co-treatment with Tumor Necrosis Factor α (TNFα) and actinomycin D (TNFα/ActD) triggered strong enhancement of extrinsic apoptosis in αmiR19b cells (Fig.1g).

### PPP2R5E subunit of PP2A is a target of miR-19b

We previously demonstrated that attenuating *miR-19b* in non-small cell lung cancer (NSCLC) cells reduced phosphorylation at serine/threonine residues by targeting serine/threonine (S/T) phosphatase PP2A subunit, PPP2R5E ^16^. This attenuation also led to increased S/T phosphatase activity (Fig.2a) and elevated *PPP2R5E* mRNA levels in GBM cells (Fig.2b). Additionally, luciferase reporter assays, using a construct containing the *miR-19b* target site (TS) of *PPP2R5E*, resulted in enhanced luciferase activity in αmiR19b compared to αsrc group, while a construct in which the *miR-19b* target site of *PPP2R5E* was mutated (mTS) did not yield to enhanced luciferase activity (Fig.2c). Attenuating *miR-19b* also resulted in reduced phospho-AKT (pAkt) consistent with PPP2R5E reactivation, but phospho-ERK (pErk) levels were surprisingly enhanced in both U87MG and GSC3 cells (Fig.2d). We found that Cellular Inhibitor of PP2A (CIP2A), an endogenous inhibitor of PP2A, was upregulated in αmiR19b cells and downregulated in pmiR19b cells (Fig.2d). Notably, the mechanism of CIP2A regulation by *miR-19b* is unknown, although it is well established that CIP2A induction contributes to Erk phosphorylation^18^.

**Fig. 2.**
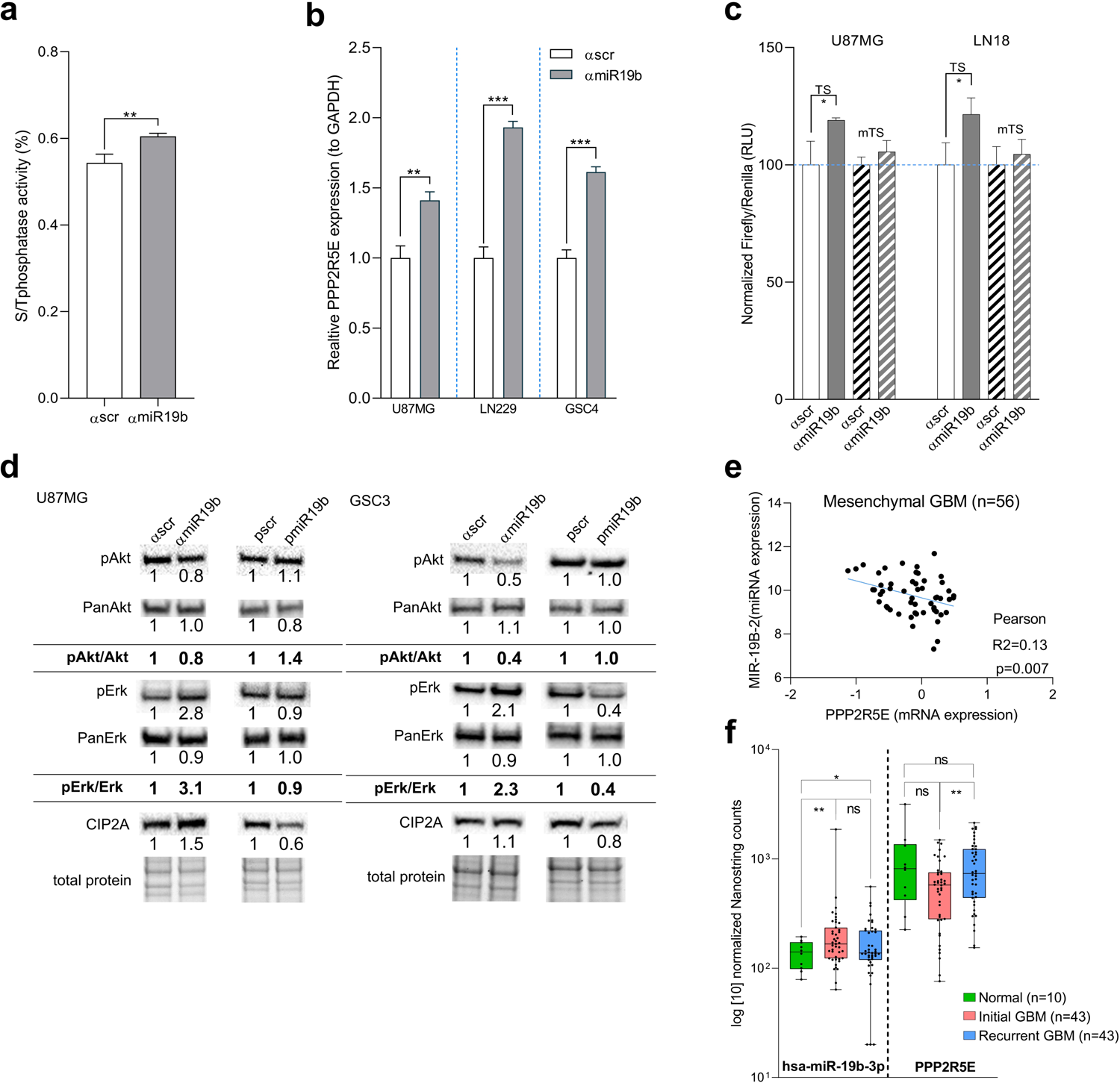
*MiR-19b* targets PP2A regulatory subunit PPP2R5E. **a** S/T phosphatase activity assay of αmiR19b U87MG cells (n=5). **b** *PPP2R5E* expression relative to *GAPDH* in αmiR19b cells by RTqPCR (n=4). **c** Luciferase activity assay of a reporter construct containing a 3’-untranslated region encompassing the *miR-19b-3p* target site of *PPP2R5E* (TS) or mutations in the *miR-19b* target site (mTS) (n=4). RLU, Relative Luminescence Unit. **d** Western blot analysis of αmiR19b and pmiR19b cells relative to control. Signal intensity of antibodies was normalized to total protein. **e** Inverse Pearson correlation of *miR-19b* and *PPP2R5E* mRNA in the mesenchymal subtype (n=56) of GBM-TCGA cases. **f** *PPP2R5E* and *miR-19b* mRNA expression in recurrent GBM. Statistics: Wilcoxon matched-paired signed rank test.

We next assessed expression of *miR-19b* and *PPP2R5E* mRNA in the TCGA-GBM public dataset (n=206). Interestingly, expression of *PPP2R5E* was inversely correlated with the expression of *miR-19b* in the mesenchymal molecular subtype (Fig.2e), but not in the classical, proneural, or neural subtypes of GBM (suppl.Fig.3a). Consistent with this finding, patients with low expression of *PPP2R5E* (z-score<-2) were more frequent in the mesenchymal (57%) and neural (52%) subtypes compared to classical (26%) and proneural (39%) subtypes, respectively (suppl.Fig.3b).

To further validate these results, we analyzed an in-house “longitudinal” cohort (n=43) consisting of matched initial (therapy-naïve) and recurrent tumors, as well as normal adjacent brain tissues ^10^. While *miR-19b* expression was significantly higher in initial tumors compared to normal tissue (Fig.2f), there was no inverse correlation between *miR-19b* and *PPP2R5E* expression in initial and recurrent tumors (Pearson R^2^=0.003). Therefore, although *miR-19b*/PPP2R5E regulatory axis plays an important role in intrinsic TMZ resistance, it does not appear to be involved in GBM recurrence.

### MiR-19b confers TMZ resistance by targeting PPP2R5E

We next investigated whether knocking down *PPP2R5E* (shPPP2R5E) phenocopies *miR-19b* overexpression in relation to TMZ resistance. Indeed, sh*PPP2R5E* enhanced cell viability (Fig.3a and suppl Fig.4a) and clonogenic growth (Fig.3b) in the presence of TMZ. On the other hand, knocking down PP2A catalytic subunit *PPP2CA* did not alter TMZ sensitivity (Fig.3a,b), despite reducing S/T phosphatase activity to an extent comparable to that of the *PPP2R5E* knockdown (suppl.Fig.4b). This implies that PPP2R5E is the essential component of PP2A complexes responsible for TMZ response. Consistent with the results of pmiR19b cells, TMZ-induced G2/M arrest was enhanced in shPPP2R5E cells (Fig.3c), and more shPPP2R5E cells could escape TMZ-induced cell cycle arrest compared to sh(ctrl), while shPPP2CA had no effect (Fig.3d). Enhanced TMZ resistance in shPPP2R5E cells was not due to higher levels of pErk and pAkt, known mediators of resistance, since these phosphoproteins were also upregulated in shPPP2CA cells (Fig.3e). Gene set enrichment analysis of shPPP2R5E cells revealed enrichment of EGFR signaling (ERBB2, ERBB4, PI3K, AKT) and p53 pathways, which may potentially contribute to TMZ resistance (suppl.Fig.4c).

**Fig. 3.**
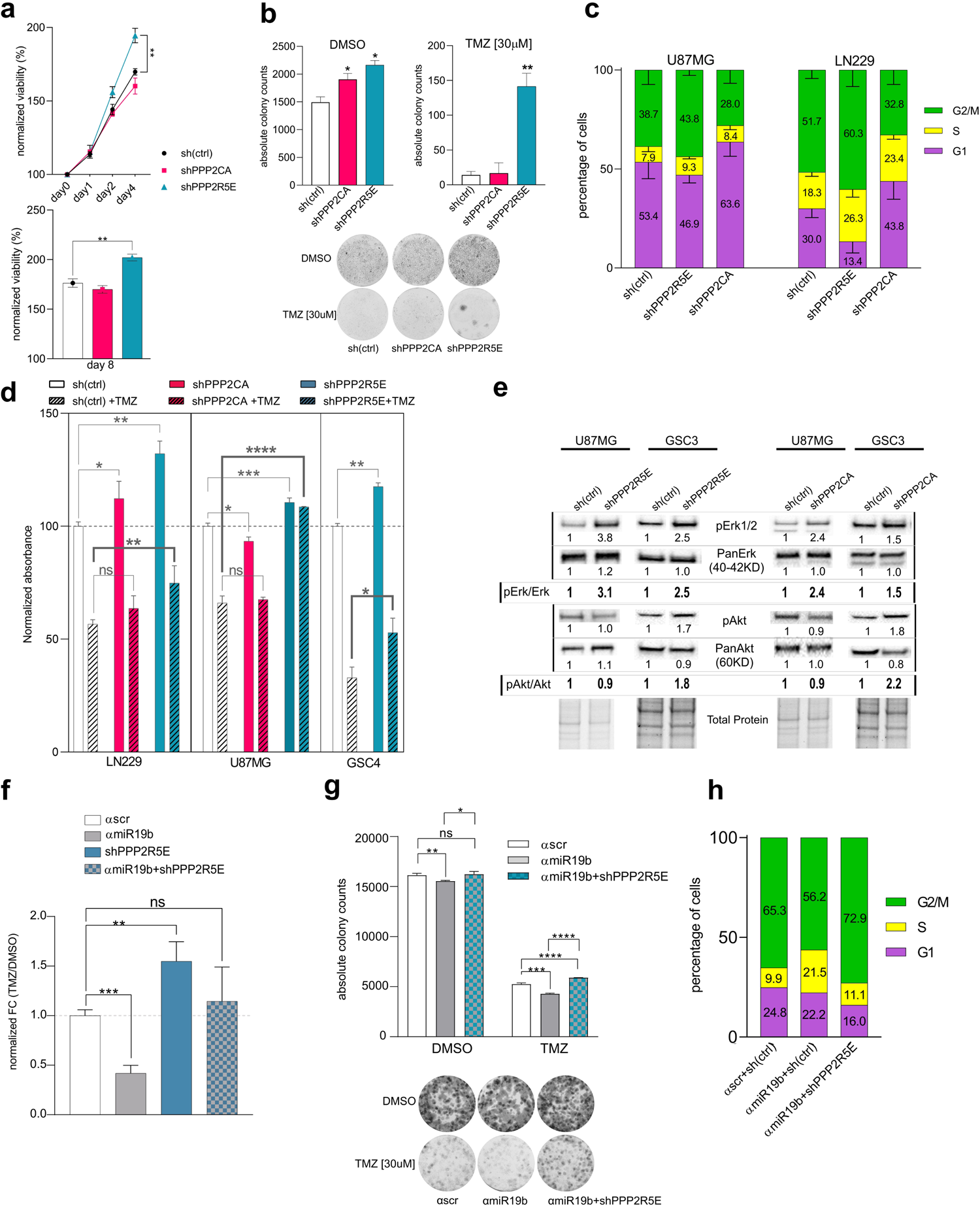
*MiR-19b*/PPP2R5E axis is implicated in TMZ resistance. **a** Cell viability by alamarBlue of shPPP2R5E U87MG cells in the presence of DMSO for 4 days (upper) or TMZ for 8 days (lower) (n=3). **b** Colony formation assay of shPPP2R5E and shPPP2CA U87MG cells in presence of continuous treatment with 30μM TMZ or DMSO control (n=3). **c** Cell cycle position following treatment with TMZ for 48h (mean±SD, n=3). **d** BrdU incorporation assay (n=4). **e** Western blot analysis. pAkt/Akt and pErk/Erk indicate the signal ratio of phospho-specific antibody and pan-antibody. **f** Cell viability by alamarBlue in presence of TMZ, relative to the DMSO control. **g** Colony formation assay (n=3) and **h** cell cycle distribution (n=1) of ɑmiR19b/shPPP2R5E co-transduced LN229 cells following TMZ treatment.

To unequivocally assess if *miR-19b* confers TMZ resistance by targeting PPP2R5E, LN229 cells were co-transduced with αmiR19b and shPPP2R5E (αmiR19b+shPPP2R5E) and treated with TMZ. αmiR19b conferred increased sensitivity whereas shPPP2R5E conferred increased resistance, but αmiR19b+shPPP2R5E cells attenuated the effects elicited by αmiR19b or shPPP2R5E alone (Fig.3f). These results were also confirmed by colony formation assay (Fig.3g). In addition, decreased TMZ-induced G2/M arrest (Fig.3h) elicited by αmiR19b was rescued in αmiR19b/shPPP2R5E cells. Hence, TMZ resistance is induced at least in part by the *miR-19b*/PPP2R5E axis.

### PP2A reactivation by other means also leads to TMZ sensitivity

We hypothesized that enhancing PP2A activity by knocking down PP2A inhibiting proteins (PAIP) ^19^ could increase TMZ sensitivity. Interestingly, in our cohort of recurrent GBM, we found that PAIP genes *CIP2A* and *PHAP1* were upregulated in recurrent tumors compared to matched initial tumors (Fig.4a), which could contribute to enhanced TMZ resistance in the recurrent tumor. To investigate this further, we generated PAIP knockdown cell lines using two different shRNAs constructs (suppl Table2, and suppl.Fig.5a-b). As expected, shPHAP1, shCIP2A#1 and shPME1 cells were more sensitive to TMZ compared to sh(ctrl), as shown by clonogenic growth assay (Fig.4b).

**Fig. 4.**
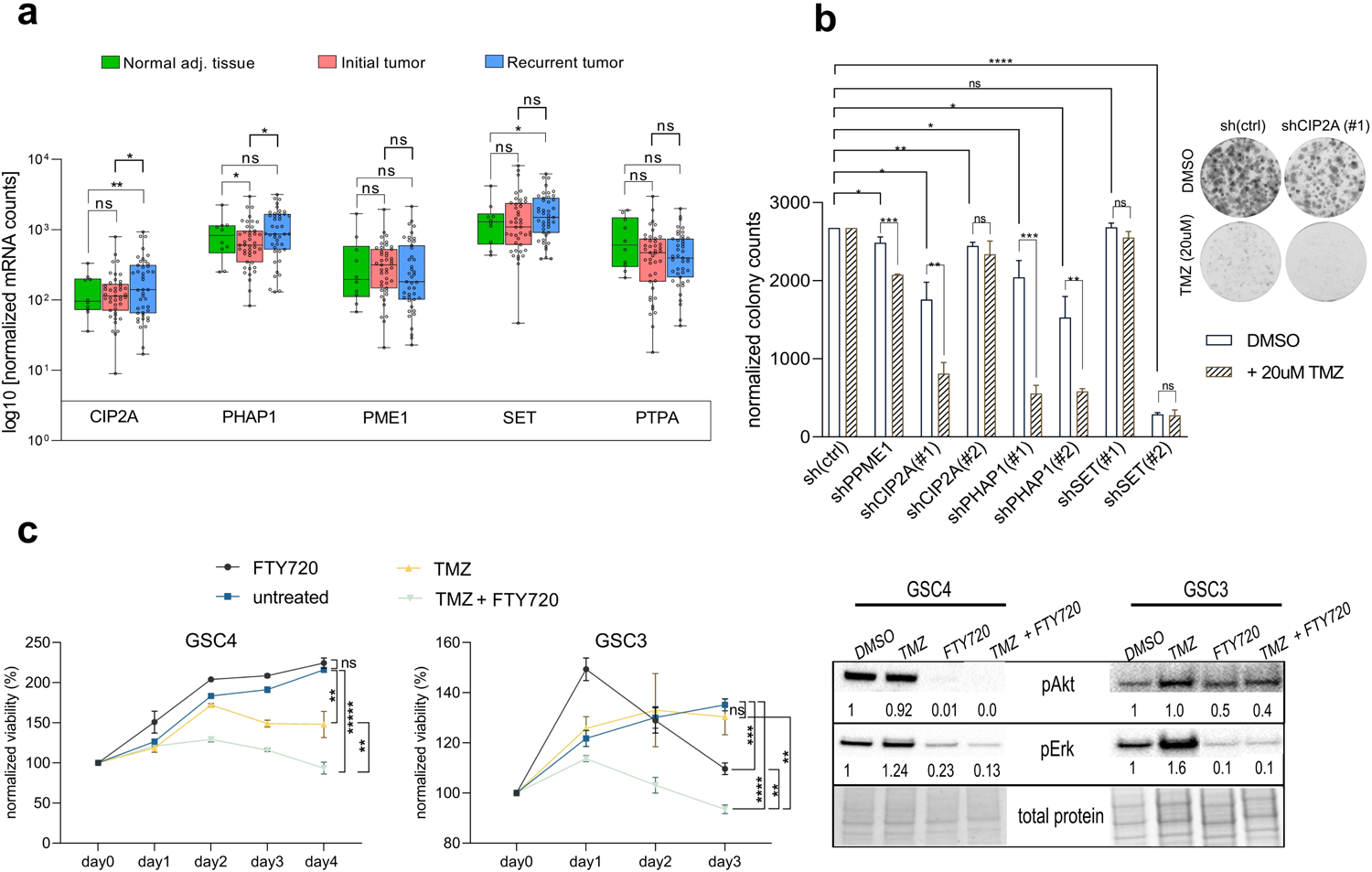
PP2A activation by means of attenuation of PP2A inhibiting proteins (PAIPs) or FTY720 treatment enhances TMZ sensitivity. **a** Expression of PAIPs in normal brain tissues and patient matched initial and recurrent GBM. **b** Colony formation assay of PAIP knockdown U87MG cells in presence of 20uM TMZ (n=3). Right: representative image of shCIP2A knockdown in a separated experiment (LN229). **c** AlamarBlue viability assay (left) and Western blot analysis for pAkt and pErk (right) performed 3-4 days post treatment of GSCs with FTY720, TMZ or combination. GSCs were treated with FTY720 at IC50.

We next aimed to activate PP2A pharmacologically using FTY720 (fingolimod), a small PP2A activator used for the treatment of multiple sclerosis ^20^. FTY720 treatment resulted in reduced pAkt and pErk levels with fast kinetics (suppl.Fig.5c). In GSCs, the combination treatment with TMZ and FTY720 showed more potent cytotoxic effects than the individual treatments (Fig.4c and suppl.Fig.5d). Therefore, reactivating PP2A through means other than *miR-19b* attenuation also potentiates the cytotoxic effect of TMZ.

### PP2A activation induces DNA damage

γH2AX foci are induced by DNA DSBs, and removed by the DNA repair machinery in a timely manner ^21^. TMZ-induced γH2AX foci formation and expression was strongly enhanced in αmiR19b LN229 (Fig.5a and suppl.Fig.6a) and U87MG cells (suppl.Fig.6b), whereas αmiR19b/shPPP2R5E cells showed attenuated foci formation (Fig.5a). Similar results, although at lower foci counts, were observed in DMSO-treated LN229 (Fig.5a), U87MG (suppl.Fig.6b, left) and GSC4 (suppl.Fig.6c) cells. To assess γH2AX foci clearance, LN229 cells were pulse-treated with TMZ for 5 hours and foci disappearance was measured during the recovery period after wash-out of TMZ. Comparable kinetics of γH2AX foci formation and clearance were observed in αscr, αmiR19b and αmiR19b+shPPP2R5E (Fig.5b), suggesting that enhanced γH2AX foci elicited by αmiR19b was not due to delayed kinetics of foci clearance. αmiR19b cells showed enlarged cellular and nuclear morphologies with a granular texture reminiscent of senescent cells (Fig.5a, brightfield channel). Indeed, αmiR19b cells expressed elevated β-galactosidase (SA-β-gal) levels compared to αscr at steady-state, and this phenotype was reverted in αmiR19b/shPPP2R5E cells. TMZ treatment strongly induced SA-β-gal expression, but again higher levels were obtained in αmiR19b compared to αscr and αmiR19b/shPPP2R5E cells (Fig.5c). In addition, TMZ-induced phosphorylation of CHK1 and CHK2 were enhanced in αmiR19b cells but reduced in shPPP2R5E cells. Conversely, FTY720 treatment enhanced TMZ-induced phosphorylation of CHK1 and CHK2, which was again reversed in shPPP2R5E cells (Fig.5d), confirming that PP2A complexes containing PPP2R5E sensitize GSCs for TMZ response. Consistent with this finding, TMZ-induced phosphorylation of CHK1 and CHK2 and γH2AX expression were enhanced in shPAIP cells (Fig.5e).

**Fig. 5.**
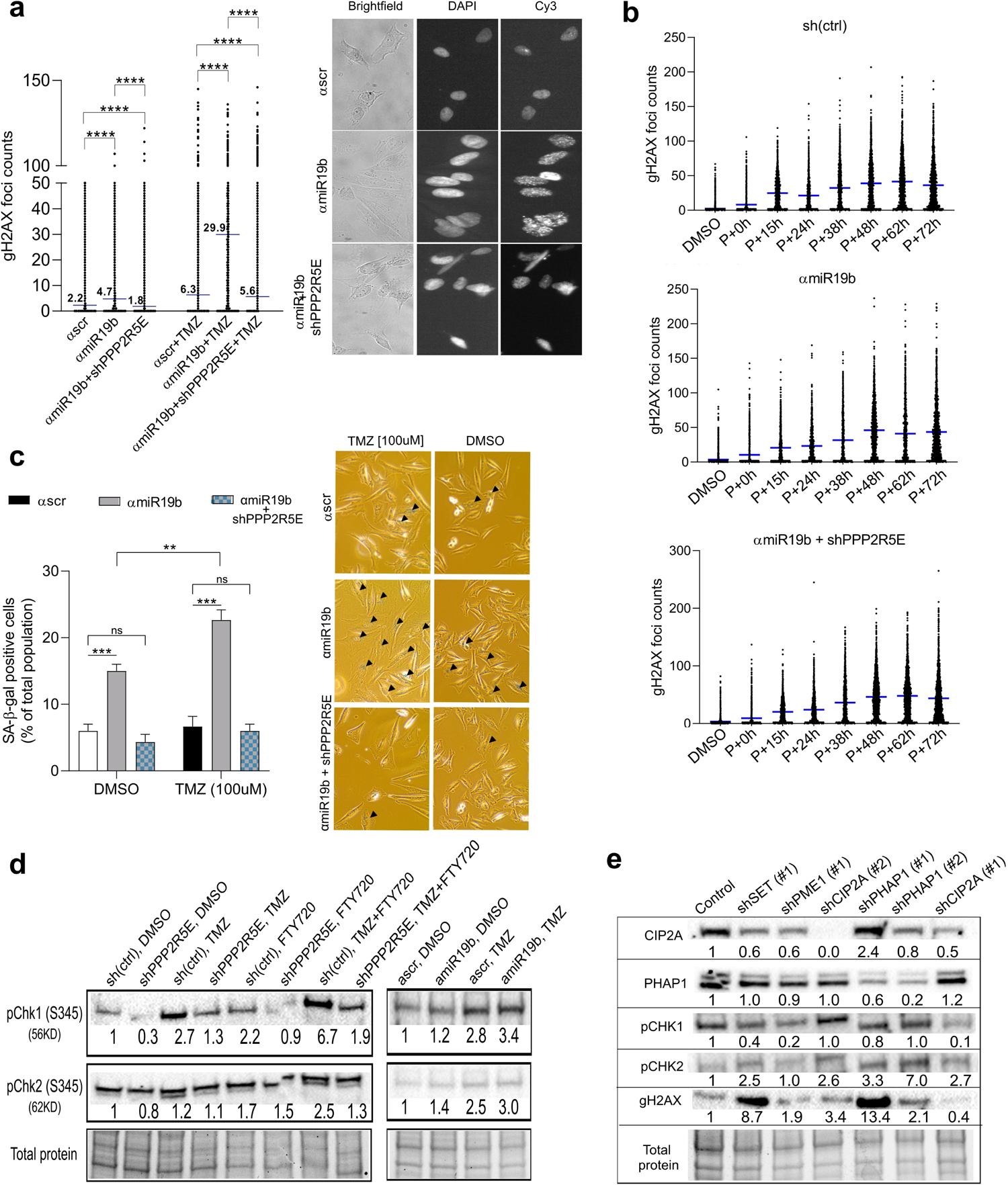
DNA damage and senescence induced by *miR-19b* attenuation is enhanced by TMZ treatment. **a** γH2AX foci counts of DMSO or TMZ-treated LN229 cells (left) and representative images (60X magnification) in bright field, DAPI (nuclei) and Cy3 (γH2AX foci). Foci were quantified using CellProfiler. The mean (n>10000) from two biological replicates is indicated by a horizontal blue line and foldchange is indicated by number **b** Kinetics of γH2AX foci clearance. LN229 sh(ctrl) (top), ɑmiR19b (middle) and ɑmiR19b+shPPP2R5E (bottom) cells were pulse-treated with 100μM TMZ for 5h (P) and chased in the absence of TMZ (drug washout/holiday) for time periods indicated in the figure. Foci were quantified using CellProfiler. Mean is indicated as horizontal blue lines and foldchange is indicated by number. **c** Percentage of senescence-associated β-galactosidase-positive DMSO or TMZ treated LN229 cells (n=4) (left) and representative images in bright field (right). Arrowheads indicate SA-β-gal**-**positive cells. **d,e** Western blot of checkpoint proteins of GSC3 cells transduced with shPPP2R5E, ɑmiR19b, FTY720, or combinations (**d)** and U87MG cells transduced with shPAIP (**e)** following treatment with TMZ for 2 days.

Taken together, these results indicate that PP2A activation by FT720 treatment or by means of *miR-19b* or PAIP attenuation potentiate DNA damages induced by TMZ treatment (Fig.6).

**Fig. 6.**
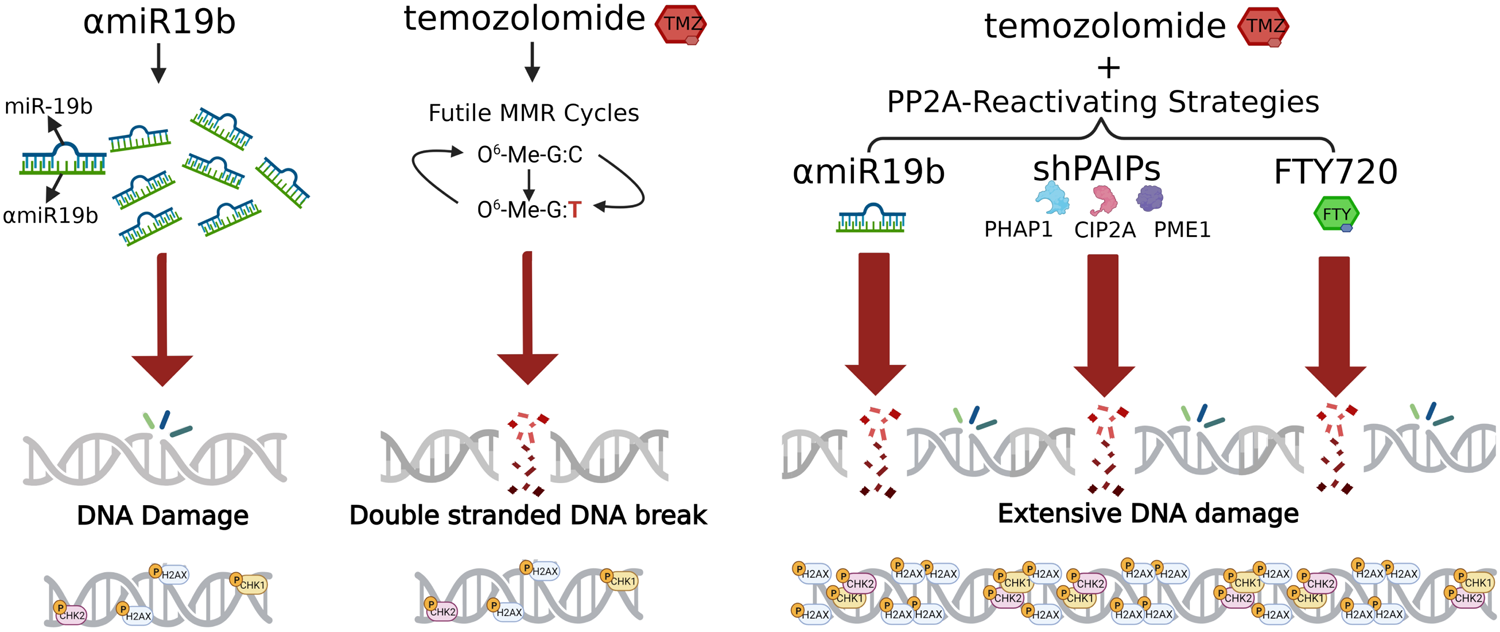
Model of combined action of DNA damage elicited by TMZ treatment and PPP2R5E reactivation. *MiR-19b* attenuation or TMZ treatment alone causes moderate DNA damage, but the combination of PP2A reactivation by means of ɑmiR19b, shPAIPs or FTY720 and TMZ treatment causes extensive DNA damage.

### PPP2R5E knockdown induces TMZ resistance in vivo

To address the physiological relevance of our findings, we next assessed TMZ response of shPPP2R5E *in vivo*. To this end, we established a patient-derived xenograft (PDX) mouse model using GSC4 cells orthotopically injected into the right brain hemisphere of nude mice. GSC4 were transduced with beetle luciferase reporter construct allowing for monitoring tumor growth using an in vivo imaging system (IVIS). Mice received five cycles of TMZ per week, which was repeated after 5 and 8 weeks (Fig.7a), much similar to the protocol used for the treatment of human patients. Unexpectedly, tumor engraftment prior to initiating therapy was compromised in mice receiving shPPP2R5E GSCs, as indicated by a smaller tumor volume 11 days post xenografting compared to the sh(ctrl) group (Fig.7b). In addition, 92% (22/24) of GSC4-sh(ctrl)-injected mice, but only 76% (16/21) of GSC4-shPPP2R5E-recipient animals developed tumors. However, once tumors had been established, shPPP2R5E developed larger tumors despite TMZ treatment (Fig.7c). Furthermore, shPPP2R5E tumors tended to be more aggressive than sh(ctrl) as indicated by Ki67 staining (Fig.7d). A tendency towards enhanced β-catenin and SMAD2/3 activity, known mediators of stem cell maintenance and proliferation following TMZ treatment ^22^, was detected in shPPP2R5E tumors, whereas expression of SOX9, a protein involved in differentiation, migration and invasion of GBM ^23^, was reduced (Fig.7d).

**Fig. 7.**
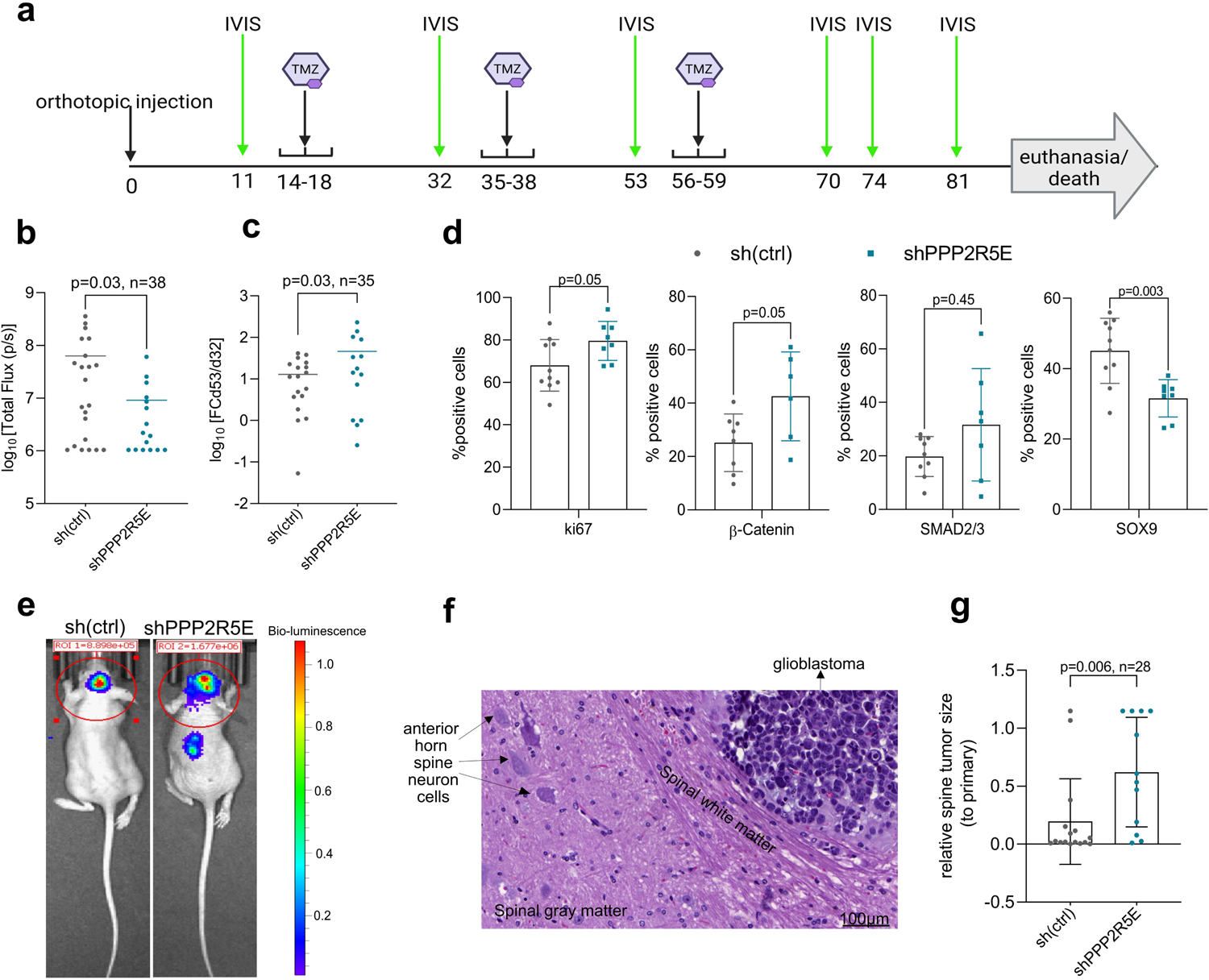
PPP2R5E attenuation contributes to TMZ resistance and distant metastasis in the GSC4 orthotopic mouse model. **a** TMZ treatment cycles and monitoring of tumor size (IVIS). **b** Tumor volume by the IVIS imaging 11 days post-xenografting prior to initiating TMZ treatment. **c** Tumor volume 52 days post-xenografting following two cycles of TMZ treatment. **d** Immunohistochemical analysis of tumors from mice euthanized at onset of neurological symptoms. **e** Example of IVIS images of sh(ctrl) and shPPP2R5E mice 74 days post-xenografting with spinal cord metastasis. **f** H&E-stained slide from spinal cord metastasis of shPPP2R5E tumor 74 days post-xenografting. **g** Tumor volume from metastatic site relative to primary tumor 32 days post-xenografting.

Of note, GSC4 tumors formed distant metastases into the left brain hemisphere and distant spinal cord (Fig.7e). GSC4 does not invade into the brain parenchyma, a finding also confirmed in distant metastases to the spinal cord (Fig.7f). While there were no significant differences in the frequency of metastases from shPPP2R5E versus sh(ctrl) tumors, the tumor volume was larger in shPPP2R5E tumors compared to sh(ctrl) at the metastatic sites (Fig.7g).

## Discussion

The mechanisms underpinning innate and acquired resistance to TMZ in GBM are still unclear. To address the role of miRNAs in TMZ response, we performed functional miRNA screens. A lentiviral construct expressing *pre-miR19b/20b* was the only construct consistently selected by TMZ in two GBM cell lines. *MiR-19b* is a critical component of oncomiR-1 involved in proliferation, apoptosis, invasion (reviewed by ^24^), and drug resistance in breast cancer ^25^ and NSCLC ^16^. It is significantly upregulated in glioma tissues and positively correlates with the glioma grade ^26^. *MiR-19b* expression is associated with poor prognosis of GBM patients treated with TMZ ^27,28^, a finding also confirmed in our study (suppl.Fig.1e). Yet, the underlying mechanisms remained so far uninvestigated.

By revealing a novel axis between *miR-19b* and PPP2R5E, our study establishes a so far unappreciated contribution of *miR-19b* to TMZ resistance of GBM. Specifically, our results show that DNA damage was induced in *miR-19b* attenuated GBM cell lines and GSCs in the absence of TMZ, and cytotoxic effects were further enhanced following TMZ treatment. In line with these findings, treating cells with FTY720, which restores PP2A activity by interacting with CIP2A and SET ^29^, or knocking down *CIP2A*, *PME1* or *PHAP1* enhanced TMZ sensitivity mediated by PPP2R5E reactivation. Hence, reactivation of PP2A complexes containing PPP2R5E potentiates the cytotoxic effect of TMZ (Fig.6). Cellular senescence, which is induced by persistent DNA damage ^30^, was also enhanced by PPP2R5E reactivation.

It is well-established that PP2A complexes dephosphorylate DNA damage response proteins CHK1, CHK2 and ATM, thereby delaying double-strand DNA break repair ^31,32^. However, we have shown that attenuation of *miR-19b* did not alter the kinetics of resolving TMZ-induced γH2AX foci. Instead, *miR-19b* attenuation caused persistent DNA damage even in the absence of TMZ. Conversely, overexpression of *miR-19b* (or *PPP2R5E* repression) resulted in less DNA damage. The mechanism of enhanced DNA damage elicited by PPP2R5E reactivation is unknown. PP2A induction is associated with enhanced senescence in melanoma ^33^, KRAS-driven lung cancer ^34^ and GBM ^35^, but the underlying mechanisms were not investigated. Phosphoproteomics and transcriptomics analysis revealed a role of the PP2A holoenzyme in inducing metabolic enzymes and NRF2-mediated oxidative stress response pathways ^36^, which may lead to elevated ROS production and enhanced DNA damage. Mitochondrial dysfunction, which gives rise to elevated ROS production, is a hallmark of cellular senescence ^37^. Thus, an alternative explanation could be that DNA damage is a consequence of PPP2R5E-induced senescence. However, it remains to be shown if PP2A reactivation is associated with enhanced ROS production in our experimental system.

Our *in vitro* findings, which showed that *PPP2R5E* repression was associated with enhanced TMZ resistance, were confirmed using an orthotopic GSC mouse model of GBM. Unexpectedly, tumor engraftment was challenged in shPPP2R5E GSC4 recipient mice, suggesting that establishment of the GBM niche or adaptation to the murine microenvironment might be disturbed by PPP2R5E repression. Moreover, GSC4 metastasis to the spinal cord, is not uncommon as up to 2% of intracranial and 25% of supratentorial GBM can disseminate to other locations, including the spinal cord ^38^. Notably, attenuation of *PPP2R5E* did not affect dissemination but had an impact on the TMZ response at the metastatic site.

Our results suggest that inhibiting *miR-19b* and subsequent reactivating PPP2R5E could be a therapeutic approach to sensitize tumors to TMZ. However, FTY720 or other PAIP interfering drugs are associated with more severe side effects, as they affect multiple PP2A complexes. The reactivation of PPP2R5E in recurrent tumors appears to be mediated by the upregulation of *CIP2A* and *PHAP1*. Therefore, although *miR-19b* is not directly involved in recurrence, αmiR19b therapy may still be effective for patients with recurrent GBM.

Several miRNA-based therapies have entered clinical phase I and II trials for the treatment of various types of cancer and other diseases (reviewed by ^39^). Although none of these clinical trials are related to the brain, the development of new carriers for the delivery of RNA therapeutics through the blood brain barrier indicates great potential of miRNA-based therapies also for CNS tumors ^40^.

## Conclusions

Our results reveal a previously unrecognized role of *miR-19b* in promoting resistance to TMZ by interfering with PPP2R5E-mediated DNA damage. Attenuation of *miR-19b*, knocking down PP2A inhibitory proteins, or treatment with the PP2A activating drug FTY720 all potentiate the cytotoxic effect of TMZ. These novel findings suggest that miRNA-based therapies targeting the *miR-19b*/PPP2R5E axis hold promise for enhancing treatment efficacy.

## Availability of data and materials

All data needed to evaluate the conclusions in the paper are present in the paper and/or the Supplementary Materials. Additional data related to this paper may be requested from the authors.

## Declarations

### Ethics approval and consent to participate

The study was approved by the Cantonal Ethics Commission of the Canton of Bern (KEK-BE:2018-00663), which waived the requirement for written informed consent. Animal experiments were performed in accordance with the Swiss Federal regulations and were previously approved by the Veterinary Office of the Canton of Bern (number BE68/2021).

### Consent for publication

All authors have agreed to publish this manuscript.

## List of abbreviations

BrdU: 5-Bromo-2′-Deoxyuridine
CIP2A: Cellular Inhibitor of PP2A
DSBs: double stranded breaks
GBM: Glioblastoma, IDH wildtype
GSCs: glioblastoma stem cells
H2AX: H2A.X variant histone
IVIS: *In Vivo* Imaging System
MGMT: O^6^-MeG DNA methyl transferase
miRNAs: microRNAs
*miR-19b*: *hsa-miR-19b-3p*
PAIPs: PP2A inhibiting Proteins
PDX: patient-derived xenograft
pre-miR: microRNA precursor
RTqPCR: Reverse Transcription quantitative real-time PCR
TMZ: temozolomide
TNFα: Tumor Necrosis Factor α

## Acknowledgements

We thank Eugenio Zoni and Marianna Kruithof-de Julio, DBMR, University of Bern, for providing constructs and assistance with the IVIS instrument, Mario Tschan, Institute of Tissue Medicine and Pathology, University of Bern, and Ren-Wang Peng, DBMR, University of Bern, for providing antibodies. We are grateful to Vivian P. Vu (Institute of Tissue Medicine and Pathology) for assistance with the *in vivo* experiments and Cornelia Schlup for assistance with the lentiviral screens. The Microscopy Imaging Center (MIC), University of Bern, Switzerland is acknowledged for granting access to the Incell 2000 instrument.

## Author contributions

E.K., S.H. and E.V. designed the study and the experiments. S.H., U.B. and N.M-W. performed the microRNA screens. E.K. performed functional in vitro experiments. C.T. and B.F. performed *miR-19b* target validation experiments. M.L. supervised orthotopic tumor injections. K.H., C.N. and J.P. performed PDX model experiments. E.K. and T.M. handled tumor annotations and pathology review of the mice H&Es. E.K. and T.M. perfomed immunohistochemical analyses. E.K., M.C.S., T.M.M, P.K., A.P. and E.V. developed the concept. M.C.S generated the CellProfiler pipeline. P.S. provided the clinical data. P.Z. provided the GSCs and the protocols of handling. E.K performed the transcriptomics and bioinformatics of the Nanostring and TCGA data and performed the GO analysis. E.K. wrote the initial draft of the manuscript and generated all the figures. E.V. acquired funding and wrote the final draft of the manuscript.

## References

1. Louis, D. N. et al. The 2021 WHO classification of tumors of the central nervous system: A summary. Neuro Oncol. 23, 1231–1251 (2021).

2. Knizhnik, A. V, Roos, W. P., Nikolova, T., Quiros, S. & Tomaszowski, K.-H. Survival and Death Strategies in Glioma Cells: Autophagy, Senescence and Apoptosis Triggered by a Single Type of Temozolomide-Induced DNA Damage. PLoS One 8, 556–65 (2013).

3. Hegi, M. E. et al. MGMT Gene Silencing and Benefit from Temozolomide in Glioblastoma. N Engl J Med 352, 997–1003 (2005).

4. Del Alcazar, C. R. G., Todorova, P. K., Habib, A. A., Mukherjee, B. & Burma, S. Augmented HR repair mediates acquired temozolomide resistance in glioblastoma. Mol. Cancer Res. 14, 928–940 (2016).

5. Annovazzi, L. et al. The DNA damage/repair cascade in glioblastoma cell lines after chemotherapeutic agent treatment. Int. J. Oncol. 46, 2299–2308 (2015).

6. Kitange, G. J. et al. Inhibition of histone deacetylation potentiates the evolution of acquired temozolomide resistance linked to MGMT upregulation in glioblastoma xenografts. Clin. Cancer Res. 18, 4070–4079 (2012).

7. Yamashiro K, Nakao K, Ohba Sh, H. Y. Human Glioma Cells Acquire Temozolomide Resistance After Repeated Drug Exposure Via DNA Mismatch Repair Dysfunction. Anticancer Res. 40, 1315–1323 (2020).

8. Zhang, X. et al. Acquired temozolomide resistance in MGMTlow gliomas is associated with regulation of homologous recombination repair by ROCK2. Cell Death Dis. 13, 138 (2022).

9. Barthel, F. P. et al. Longitudinal molecular trajectories of diffuse glioma in adults. Nature 576, 112–120 (2019).

10. Kashani, E. et al. Integrated longitudinal analysis of adult grade 4 diffuse gliomas with long-term relapse interval revealed upregulation of TGF-β signaling in recurrent tumors. Neuro Oncol. 16:noac265, (2022).

11. Peraza-Vega, R. I., Valverde, M. & Rojas, E. Interactions between miRNAs and Double-Strand Breaks DNA Repair Genes, Pursuing a Fine-Tuning of Repair. Int. J. Mol. Sci. 23, 3231 (2022).

12. Haemmig, S. et al. MiR-125b controls apoptosis and temozolomide resistance by targeting TNFAIP3 and NKIRAS2 in glioblastomas. Cell Death Dis. 5, e1279 (2014).

13. Luedi, M. M. et al. Dexamethasone-mediated oncogenicity in vitro and in an animal model of glioblastoma. J. Neurosurg. 129, 1446–1455 (2018).

14. Hall, M. P. et al. Click beetle luciferase mutant and near infrared naphthyl-luciferins for improved bioluminescence imaging. Nat. Commun. 9, 132 (2018).

15. Frenster J & Placantonakis D. Bioluminescent In Vivo Imaging of Orthotopic Glioblastoma Xenografts in Mice. in *Glioblastoma*, Methods and Protocols vol. 1741 191–198 (Springer New York, 2018).

16. Baumgartner, U. et al. miR-19b enhances proliferation and apoptosis resistance via the EGFR signaling pathway by targeting PP2A and BIM in non-small cell lung cancer. Mol. Cancer 17, 1–15 (2018).

17. Oh, S. J. et al. Human U87 glioblastoma cells with stemness features display enhanced sensitivity to natural killer cell cytotoxicity through altered expression of NKG2D ligand. Cancer Cell Int. 17, 1–9 (2017).

18. Chen, K. F. et al. Cancerous inhibitor of protein phosphatase 2A (CIP2A) is an independent prognostic marker in wild-type KRAS metastatic colorectal cancer after colorectal liver metastasectomy. BMC Cancer 15, 1–9 (2015).

19. Kashani, E. & Vassella, E. Pleiotropy of PP2A Phosphatases in Cancer with a Focus on Glioblastoma IDH Wildtype. Cancers (Basel). 14, 5227 (2022).

20. Kappos, L. et al. A Placebo-Controlled Trial of Oral Fingolimod in Relapsing Multiple Sclerosis. N Engl J Med. 362, 387–401 (2010).

21. Mariotti, L. G. et al. Use of the γ-H2AX assay to investigate DNA repair dynamics following multiple radiation exposures. PLoS One 8, 1–12 (2013).

22. Ulasov, I. V., Nandi, S., Dey, M., Sonabend, A. M. & Lesniak, M. S. Inhibition of sonic hedgehog and notch pathways enhances sensitivity of cd133+ glioma stem cells to temozolomide therapy. Mol. Med. 17, 103–112 (2011).

23. Chao, M. et al. TGF-β Signaling Promotes Glioma Progression Through Stabilizing Sox9. Front. Immunol. 11, 592080 (2021).

24. Wang, W., Zhang, A., Hao, Y., Wang, G. & Jia, Z. The emerging role of miR-19 in glioma. J Cell Mol Med. 22, 4611–4616 (2018).

25. Liang, Z., Li, Y., Huang, K., Wagar, N. & Shim, H. Regulation of miR-19 to breast cancer chemoresistance through targeting PTEN. Pharm. Res. 28, 3091–3100 (2011).

26. Jia, Z. et al. MiR-19a and miR-19b overexpression in gliomas. Pathol. Oncol. Res. 19, 847–853 (2013).

27. Schnabel, E. et al. Prognostic value of microRNA-221/2 and 17-92 families in primary glioblastoma patients treated with postoperative radiotherapy. Int. J. Mol. Sci. 22, 1–18 (2021).

28. Zhi, F. et al. Identification of 9 serum microRNAs as potential noninvasive biomarkers of human astrocytoma. Neuro. Oncol. 17, 383–391 (2015).

29. Cristóbal, I. et al. Potential anti-tumor effects of FTY720 associated with PP2A activation: a brief review. Curr. Med. Res. Opin. 32, 1137–1141 (2016).

30. D’Adda Di Fagagna, F. Living on a break: Cellular senescence as a DNA-damage response. Nat. Rev. Cancer 8, 512–522 (2008).

31. Freeman, a; Dapic, V; Monteiro, A. Chk2/Pp2a&DNAdamage. Cell cycle 9, 736–747 (2011).

32. Leung-Pineda, V., Ryan, C. E. & Piwnica-Worms, H. Phosphorylation of Chk1 by ATR Is Antagonized by a Chk1-Regulated Protein Phosphatase 2A Circuit. Mol. Cell. Biol. 26, 7529–7538 (2006).

33. Mannava, S. et al. PP2A-B56α controls oncogene-induced senescence in normal and tumor human melanocytic cells. Oncogene 31, 1484–1492 (2012).

34. Kauko, O. et al. PP2A inhibition is a druggable MEK inhibitor resistance mechanism in KRAS-mutant lung cancer cells. Sci. Transl. Med. 10, eaaq1093 (2018).

35. Cucinotta, L. et al. The Pivotal Role of Protein Phosphatase 2A (PP2A) in Brain Tumors. Int. J. Mol. Sci. 23, 15717 (2022).

36. Chen, L. et al. Protein phosphatase 2A regulates cytotoxicity and drug resistance by dephosphorylating AHR and MDR1. J. Biol. Chem. 298, 101918 (2022).

37. Shmulevich, R. & Krizhanovsky, V. Cell Senescence, DNA Damage, and Metabolism. Antioxid Redox Signal. 34, 324–334 (2021).

38. Shah, A., Redhu, R., Nadkarni, T. & Goel, A. Supratentorial glioblastoma multiforme with spinal metastases. J. Craniovertebr. Junction Spine 1, 126–129 (2010).

39. Chakraborty, C., Sharma, A. R., Sharma, G. & Lee, S. S. Therapeutic advances of miRNAs: A preclinical and clinical update. J. Adv. Res. 28, 127– 138 (2021).

40. Petrescu, G. E. D., Sabo, A. A., Torsin, L. I., Calin, G. A. & Dragomir, M. P. MicroRNA based theranostics for brain cancer: Basic principles. J. Exp. Clin. Cancer Res. 38, 231 (2019).

