## Supplementary figures and images for "Unlocking the potential of *miR-19b* in the regulation of temozolomide response in glioblastoma patients via targeting PPP2R5E, a subunit of the protein phosphatase 2A complex"

### Supplementary Figures

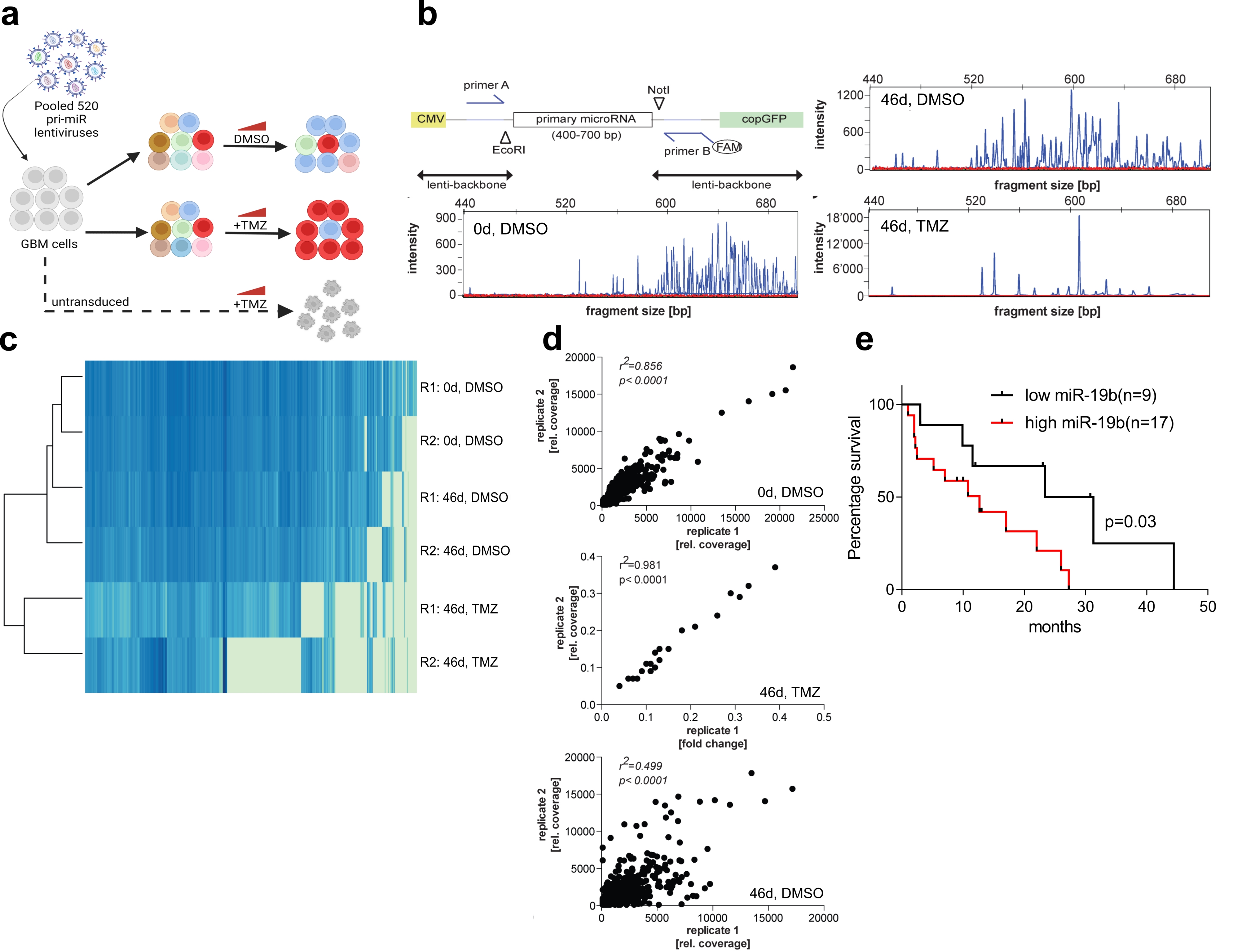

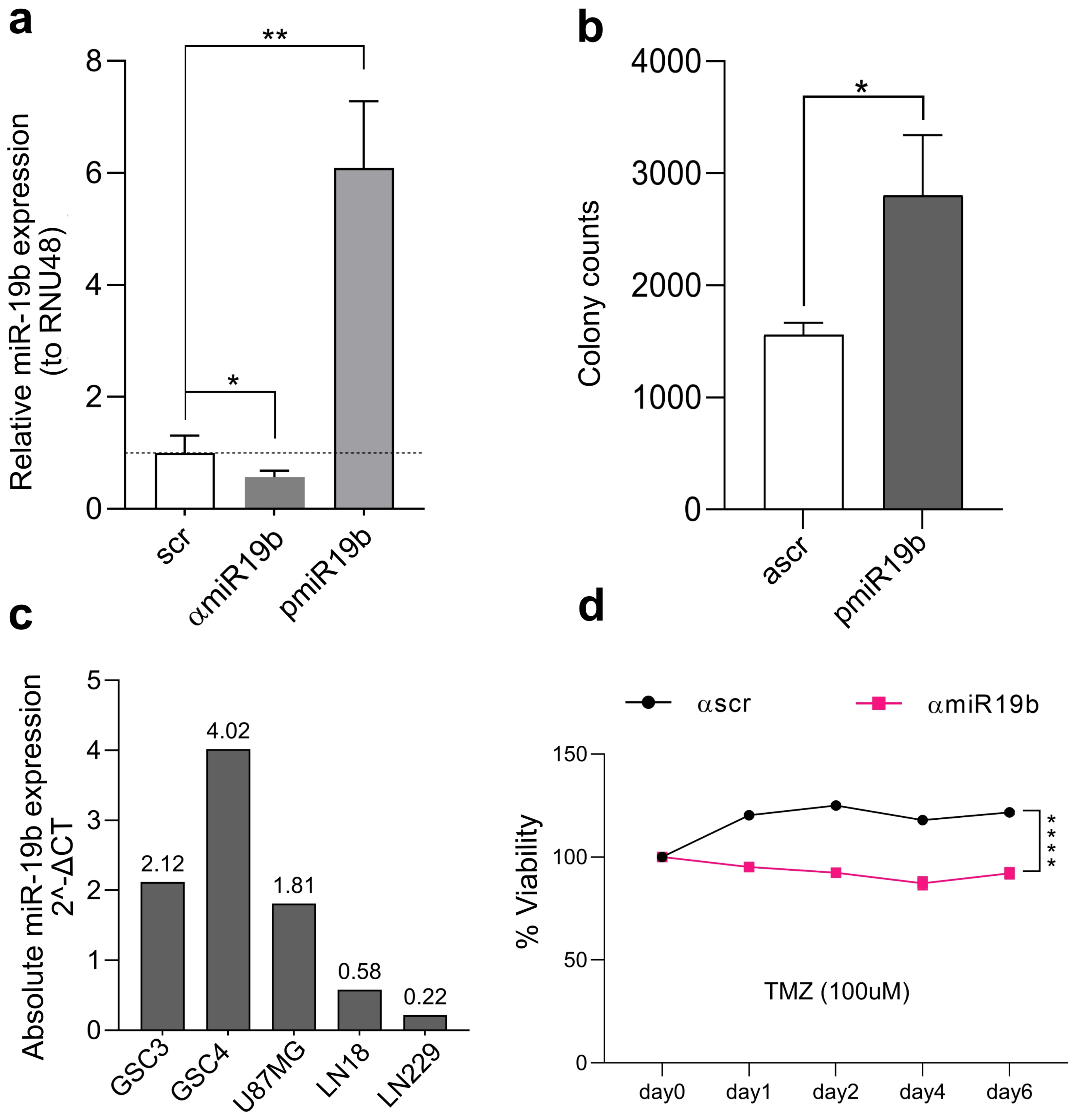

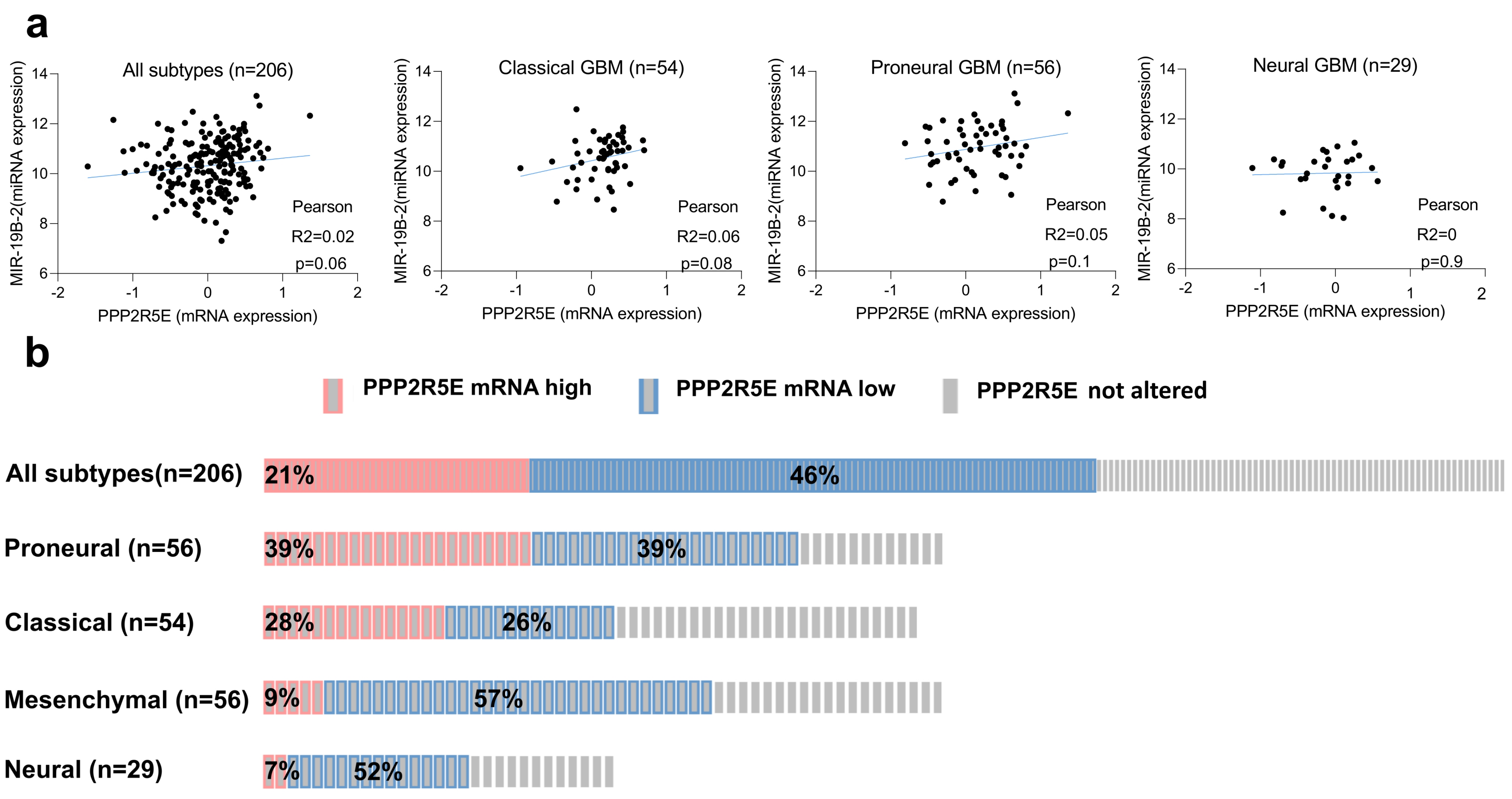

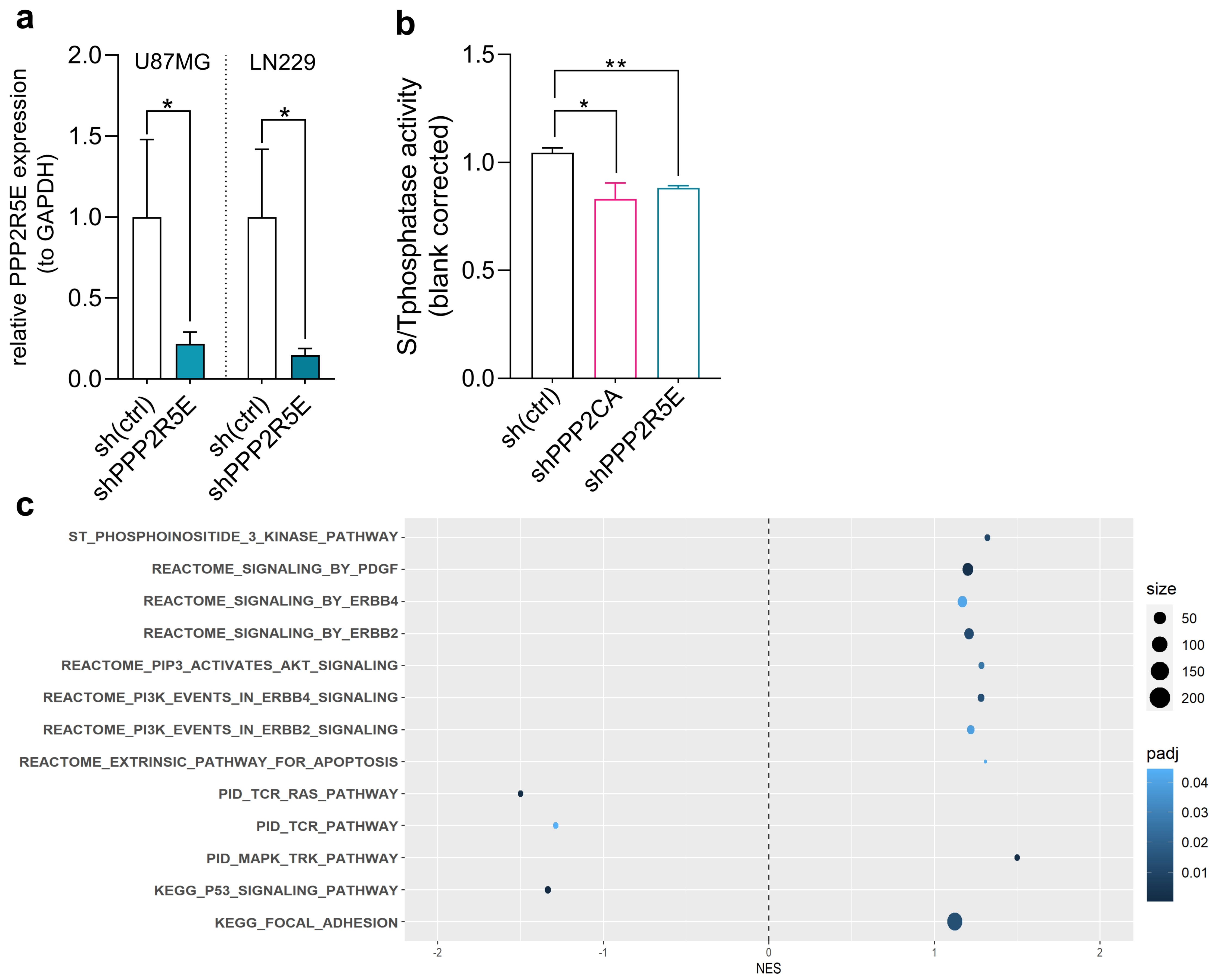

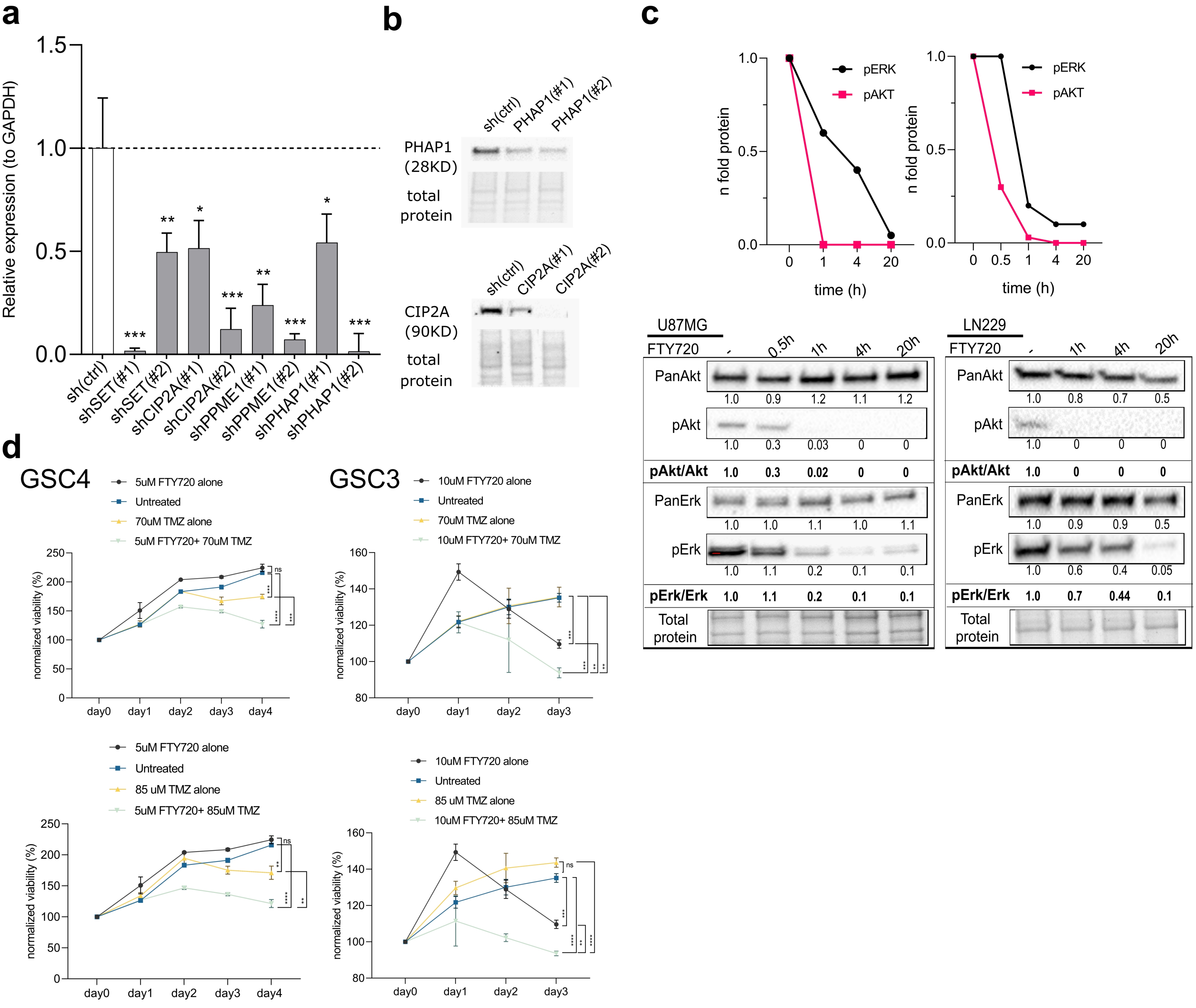

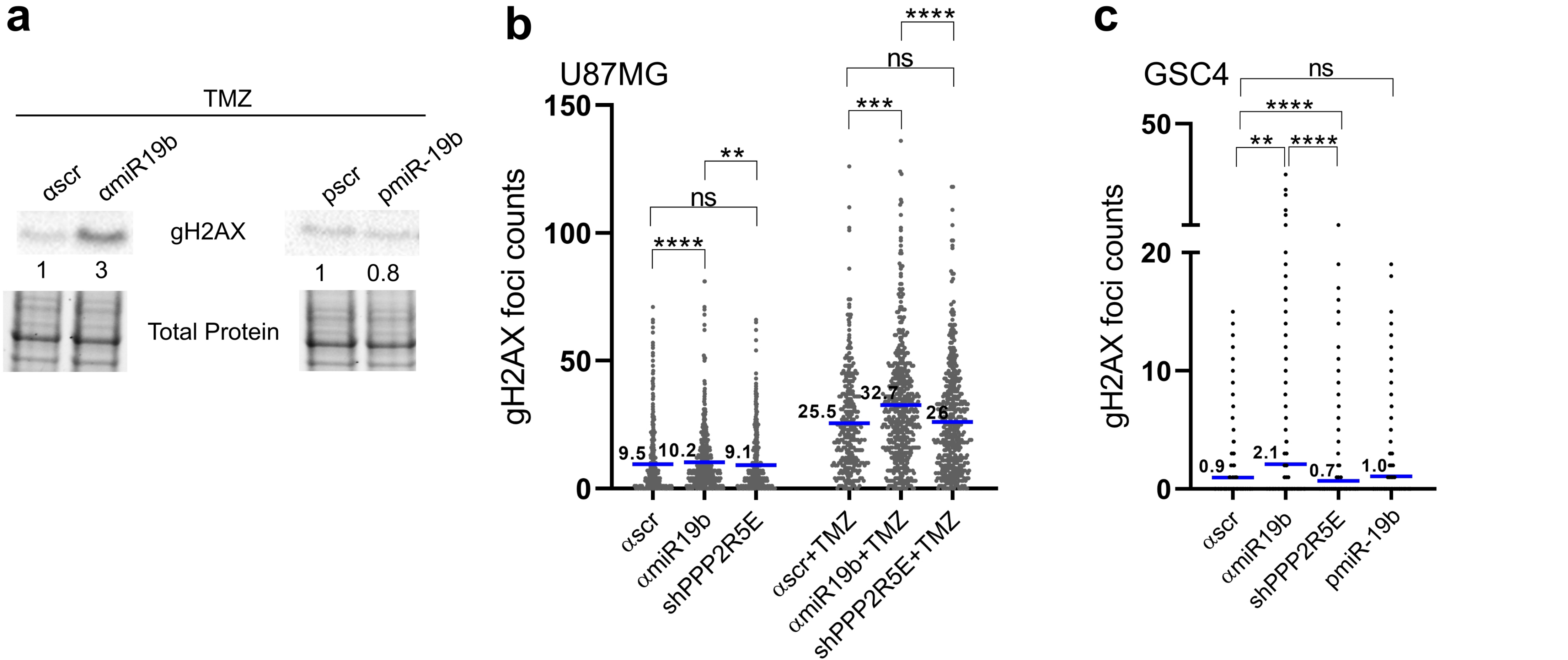
