## Supplementary Figure Legends for "Unlocking the potential of *miR-19b* in the regulation of temozolomide response in glioblastoma patients via targeting PPP2R5E, a subunit of the protein phosphatase 2A complex"

**Fig. S1 Pooled lentiviral miRNA screen.** **a** Experimental procedure for the identification of miRNAs conferring TMZ resistance. Un-transduced cells are indicated in gray. **b** PCR design for pre-miRNA fragment amplification (upper left) and fragment size distribution of U87MG cells shortly after transduction (0d, DMSO), 46d post-transduction, DMSO ctrl (46d, DMSO), and 46d post-transduction, TMZ treated (46d TMZ). **c** Heat map analysis of two independent U87MG replicates, R1 and R2. **d** Correlation of R1 and R2 for U87MG cell lines at 0d, 46d TMZ and 46d DMSO. **e** Overall survival of GBM IDH wt patients treated with TMZ in combination with radiotherapy from the discovery cohort expressing high or low levels of *miR-19b* by RT-qPCR (n=3).

**Fig. S2** **Lentiviral transduction efficiency and cellular processes affected by miR-19b.** **a** *miR-19b* expression in U87MG-transduced cells normalized to the level obtained for *RNU48* by RTqPCR (TaqMan) (n=4). **b** Clonogenic growth of U87MG cells transduced with pmiR19b and treated with 15 μM TMZ for two weeks. **c** Endogenous level of *miR-19b* in GBM stem cells and GBM cell lines by RTqPCR (n=4). **d** AlamarBlue viability assay of αmiR19b GSC4 cells relative to αscr control. Cells were treated with 100 μM TMZ for 6 days (n=4).

**Fig. S3 MiR-19b targets PP2A regulatory subunit PPP2R5E in mesenchymal GBM subtype.** **a** Pearson correlation of *miR-19b* and *PPP2R5E* mRNA in all subtypes together (n=206), proneural (n=56), classical (n=54) or neural (n=29) subtypes from GBM-TCGA dataset. Pearson correlation coefficient (R2) and p-values are indicated on each panel. **b** Oncoprint for *PPP2R5E* expression levels based on dichotomized (z-score) values in GBM subtype from TCGA-GBM dataset (n=206). Z-scores >2 are defined as high, z-scores <-2 are defined as low groups and z-scores ≤2 and ≥-2 are defined as not altered. The percentage of high and low expression groups is indicated.

**Fig. S4 PP2A subunits knockdown efficiency and deregulated pathways conferred by shPPP2R5E.** **a** *PPP2R5E* mRNA expression normalized to the level obtained for *GAPDH* by RTqPCR (n=4). **b** Ser/Thr phosphatase activity of GSC3 cells transduced with shPPP2CA or shPPP2R5E relative to sh(ctrl) (n=5). **c** Deregulated pathways in shPPP2R5E U87MG and LN229 cells. Size, number of leading genes involved in each pathway; padj, adjusted p-value; NES, normalized enrichment score.

**Fig. S5 PAIPs knockdown efficiency or FTY720 treatment and AlamarBlue viability assays in the presence of TMZ.** **a** PAIP mRNA expression in U87MG-transduced cells normalized to *GAPDH* by RTqPCR (n=4). **b** PHAP1 and CIP2A protein levels in U87MG-transduced cells by Western blot normalized to total protein. **c** Time-dependent phosphorylation of AKT and ERK following treatment with 5 μM FTY720 in U87MG and corresponding Western blots in U87MG and LN229. **d** AlamarBlue viability assay of GSCs treated with TMZ, DMSO, FTY720, or combination for 4 days. FTY720 was administered at IC50 (5uM and 10uM for GSC4 and GSC3, respectively).

**Fig. S6 γH2AX expression and foci formation is affected by PPP2R5E reactivation. a** γH2AX expression by Western blot in LN229 ɑmiR19b and pmiR19b cells treated with TMZ [100uM] for 2d. **b** γH2AX foci formation in αmiR19b and shPPP2R5E U87MG cells treated with TMZ for 2d (+TMZ) or DMSO ctrl. **c** γH2AX foci formation in non-treated αmiR19b, pmiR19b and shPPP2R5E GSC4 cells.
